# Long-chain lipids facilitate insertion of large nanoparticles into membranes of small unilamellar vesicles

**DOI:** 10.1101/2021.07.12.452073

**Authors:** Marzouq Adan, Morgenstein Lion, Carlos A. Huang-Zhu, Shimon Yudovich, Atkins Ayelet, Grupi Asaf, Reid C. Van Lehn, Weiss Shimon

## Abstract

Insertion of hydrophobic nanoparticles into phospholipid bilayers is limited to small particles that can incorporate into the hydrophobic membrane core in between the two lipid leaflets. Incorporation of nanoparticles above this size limit requires development of challenging surface engineering methodologies. In principle, increasing membrane thickness should facilitate incorporation of larger nanoparticles. Here, we explore the effect of incorporating very long phospholipids (C24:1) into small unilamellar vesicles on the membrane insertion efficiency of hydrophobic nanoparticles that are 5-13 nm in diameter. To this end, we improved an existing vesicle preparation protocol and utilized cryogenic electron microscopy imaging to examine the mode of interaction and to evaluate the insertion efficiency of membrane-inserted nanoparticles. We also perform classical, coarse-grained molecular dynamics simulations to identify changes in lipid membrane structural properties that may increase insertion efficiency. Our results indicate that long-chain lipids increase the insertion efficiency by preferentially accumulating near membrane-inserted nanoparticles to reduce the thermodynamically unfavorable disruption of the membrane.

## Introduction

In recent years, synthesis methods for the production of high-quality nanoparticles (NPs) have greatly improved, allowing for increased control over their size, shape, chemical and physical properties. This, in turn, enables the fabrication of sophisticated metallic, magnetic, dielectric, and semiconducting heterostructured NPs with desirable photophysical and chemical properties.^1–4^ This precise control allows the engineering of excited[state wavefunctions^5–7^, charge confinement, and spatiotemporal control of charge[separated states^8^. Hence, NPs (and in particular semiconducting NPs) have proved to be very useful in diverse applications such as in optoelectronics^9,10^, biological imaging^11^, sensing^12–14^, catalysis^15^, energy harvesting^16^, biomedicine, and cell surface engineering^17–20^.

While these sophisticated inorganic nanomaterials require advanced synthesis methods, their integration with biological macromolecules and machineries requires additional functionalization steps. Current approaches typically target NPs to interact with the cell membrane’s surface or to undergo cellular uptake. Less effort has been invested in the functionalization of NPs that could be targeted, incorporated into, and retained in the membrane bilayer core itself.^21^ Such membrane-inserted NPs could expand the repertoire of desired cellular functions while taking advantage of the superior inorganic materials’ properties.^22,23^ Once inserted into the membrane, such NPs could, for example, introduce orthogonal (to native signaling) ways to communicate with the cell’s interior, introduce de novo or enhance native enzymatic/catalytic activities, sense the membrane potential, or be used as antennas for light harvesting and vision restoration.^4,24,25^

The underlying obstacle for stable insertion of large NPs into lipid membranes is that, in many cases, the surfaces of as-synthesized NPs are decorated with a mixture of highly hydrophobic ligands and are insoluble in biologically-relevant aqueous media. Therefore, without any surface modification, these hydrophobic NPs must be efficiently incorporated into synthetic vesicle membranes in between the two leaflets. Since phospholipid bilayers are typically 4–5 nm thick^26,27^, successful membrane insertion has been achieved only for small NPs with sizes ≤ 5 nm^28–30^. Inclusion of larger hydrophobic NPs is thermodynamically unfavorable due to the energetic penalty associated with the protrusion of hydrophobic ligands into the surrounding polar solvent^30^ or due to the deformation of the bilayer to minimize such protrusion^31–33^. This limitation practically excludes higher-order structures and functionalities that could be beneficial in terms of signal or actuation strength.

We hypothesized that increased membrane thickness should promote the insertion of larger NPs than previously accomplished. The thickness of the phospholipid membrane can be increased by the incorporation of lipids with a long alkane chain and by the addition of cholesterol (CHOL), which aligns and stretches alkane chains into a packed and thick membrane^26^. To test this hypothesis, we prepared small unilamellar lipid vesicles incorporating 1,2-dinervonoylsn-glycero-3-phosphocholine (PC24), an unsaturated zwitterionic phospholipid with 24 carbon alkane chains. This lipid has been previously used in the preparation of multilamellar liposome (MLV) formulations with protein or peptide drugs, and in the formation of planar bilayer lipid membranes^34^. In biological systems, it is a component of lipid rafts and microdomains^35^.

Compared to typical PC18 lipids, which form membranes of 4-5 nm thickness, lipid bilayers consisting of 25% PC24 are expected to lead to a 20% increase in membrane thickness, i.e., up to 6 nm. To test our hypothesis, we evaluated the membrane-insertion efficiency of inorganic NPs in the size range of 5-13 nm in diameter into the lipid bilayer of small unilamellar vesicles (SUVs) containing various molar fractions of PC24 during the formation of lipid vesicles. Initial attempts to incorporate even small NPs into SUVs prepared by the hydration method^36^ were unsuccessful. After testing several other SUV preparation protocols for NP insertion that failed, we identified in the literature a protocol that was previously used to encapsulate hydrophobic drugs into liposomes’ membranes^37^ but never applied for membrane incorporation of nanoparticles. This approach proved to be highly efficient for membrane insertion of small NPs. A critical step in this protocol is the addition of a lipid detergent to the organic solution containing both lipids and NPs. With the addition of PC24 to the lipid mixture^34^, we found that NPs with diameters of up to ∼13 nm were successfully incorporated into SUV membranes.

Here, we present a detailed protocol for the incorporation of large NPs (up to ∼13 nm in diameter) into the lipid membrane of SUVs with optimized lipid composition, detergent, organic solvents, and NP/lipid ratio. Cryogenic electron microscopy (cryo-EM) imaging of these membrane-inserted NPs is used for the characterization of their mode of interaction with the membrane, indicating their insertion to the midplane of the membrane. To determine the role of PC24 in facilitating insertion, we further performed coarse-grained molecular dynamics simulations to interrogate lipid structural organization around membrane-embedded NPs. The simulations show that the longer PC24 lipids accumulate near membrane-inserted NPs to reduce the degree of membrane disruption and minimize exposure of hydrophobic area to solvent, thereby modulating the insertion efficiency of large NPs even without substantial changes in overall membrane thickness.

## Results

In order to insert NPs efficiently into the lipid bilayer of SUVs, we adopted a detergent dialysis-based vesicle preparation method that was previously optimized for membrane incorporation of small hydrophobic drugs^37^. This procedure was modified to allow for the insertion of NPs into SUV membranes. Two different mixtures of lipids were used for SUV preparation: a ‘native membrane’ mixture with POPC, CHOL and DOTAP (central column in Table 1), and a ‘thick membrane’ mixture with POPC, CHOL, DOTAP, and a very long-chain phospholipid (PC24). Insertion efficiency was tested for four different sizes of NPs: 5, 7, and 11 nm diameter spherical CdSe/ZnS quantum dots (QDs), and 5x13 nm cylindrical-shaped nanorods (NRs) (sizes determined by TEM, Table S1). Cryo-EM imaging was used to assess the mode of NP interaction/insertion with/into the SUVs’ membrane. As described in the Supplementary Information (SI) section, the yield of NP insertion into the membrane was evaluated by examining the cryo-EM images and counting membrane-inserted NPs, membrane-adsorbed NPs, and encapsulated NPs. We excluded all NPs that were either interacting with the carbon grid or aggregated. Multiple experiments on different days were conducted for each condition (NP size/type and ‘native’ or ‘thick’ membrane). Preparations were repeated for each condition, and multiple field-of-views (FOVs) were collected from each grid. Insertion efficiency was defined as the ratio between the number of membrane-inserted NPs and the total number of NPs in a frame. Results from all analyzed frames per condition were averaged. We note that the surfaces of the QDs and NRs are coated with hydrophobic ligands as prepared, facilitating their favorable interaction with the membrane core.

**Table 1.** Five different lipid compositions used for the NR insertion studies with [PC24] increasing from 0 mol% (sample 1) to 100 mol% (sample 5).

| Composition<br>Sample | PC24<br>(mol %) | POPC<br>(mol %) | CHOL<br>(mol %) | DOTAP<br>(mol %) |
| --- | --- | --- | --- | --- |
| 1 | 0 | 70 | 20 | 10 |
| 2 | 25 | 51 | 14 | 10 |
| 3 | 50 | 31 | 9 | 10 |
| 4 | 75 | 12 | 3 | 10 |
| 5 | 100 | 0 | 0 | 0 |

In order to find the optimal concentration of PC24 to add to the ‘thick membrane’ mixture, we first tested five preparations of SUVs using different relative concentrations of PC24 and evaluated NR insertion efficiency as a function of PC24 concentration (Table S2). NR-inserted SUVs were prepared as described in the SI, with NR concentrations carefully maintained for all 5 preparations. Each preparation was imaged by cryo-EM and subsequently analyzed to assess the mode of NR-membrane interaction. Figure 1 shows NR membrane insertion efficiency as a function of the long-chained lipid concentration [PC24] in the lipid mixture. The optimal concentration of [PC24] is 25 mol% of the total lipid molar concentration, resulting in an insertion efficiency that is twice as high as the insertion efficiency of SUVs prepared with the native membrane (2 out of 7 particles in the native membrane compared to 5 out of 10 particles in the thick membrane). Higher or lower [PC24] showed a decreased insertion efficiency compared to [PC24] = 25 mol%. Although [PC24] = 100 mol% also resulted in good NR insertion, which was comparable to [PC24] = 25 mol%, the morphology of these SUVs was not spherical (Figure S2). The non-spherical, polygonal SUV shapes can be attributed to the membrane’s gel phase behavior, since PC24 has a transition temperature of 26°C^38–40^, which is higher than the transition temperature of POPC (-2°C). In addition, at [PC24] = 50 mol%, SUV formation remarkably decreased and no NPs were observed upon cryoEM imaging (data not shown).

**Figure 1.**
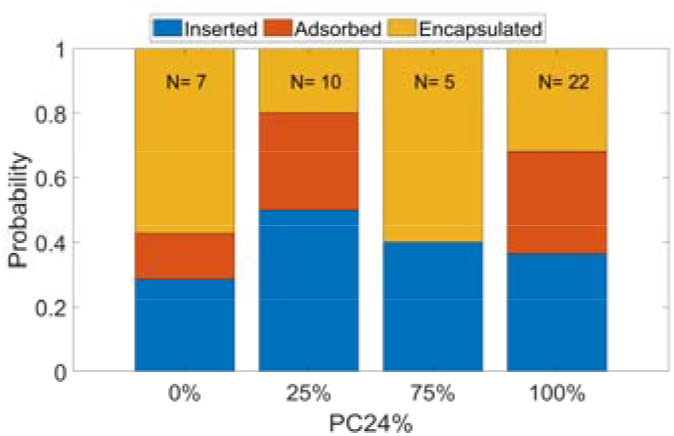
NR membrane insertion efficiency as a function of PC24 lipid molar percentage in the SUV membrane composition. N is defined as membrane interacting monodisperse NPs. SUVs with 100% PC24 exhibited non-round, distorted shapes, as shown in Figure S2.

Figure 2 shows cryo-EM images of selected examples of successful insertion of the different size/type of NPs into SUVs created from the ‘thick’ membrane mixture. The morphologies of the SUVs at different PC24 concentrations are shown in SI Figure S2 and raw images are deposited^40^. The blue and red bars on the right columns indicate the fractions of inserted and non-inserted particles, respectively, out of all counted particles, excluding aggregates and carbon-interacting particles.

**Figure 2.**
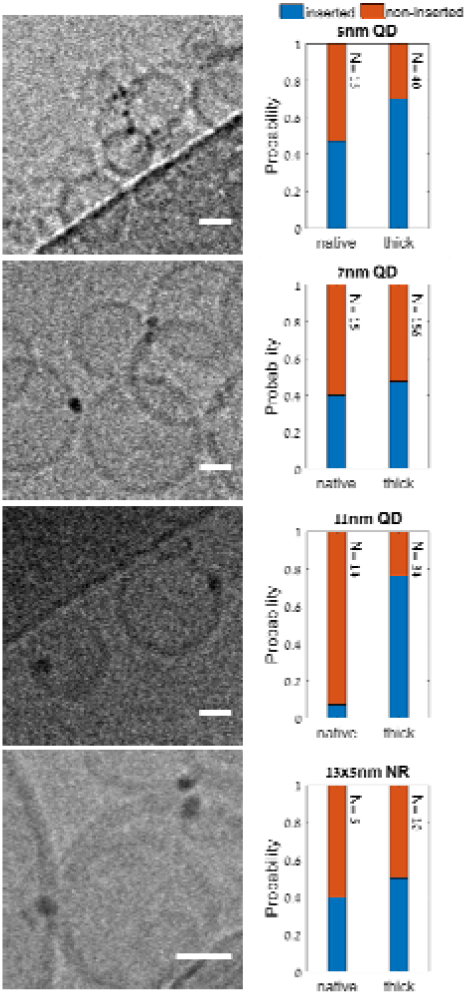
Incorporation of NPs into native and thick SUV membranes. Left column: cryo-EM images showing examples for insertion of 5 nm, 7 nm, 11 nm QDs, and 5x13 nm NRs (from top to bottom) into thick SUV membranes. Right column: corresponding insertion efficiencies of NPs in native and thick membranes. Lipid compositions for native and thick membranes are described in Table 2. Scale bars: 20 nm.

As expected, insertion efficiency was inversely proportional to the NP size for native membranes due to the NP size, whereas the long lipid addition had a significant effect on the larger particle as a result of the increase of the membrane thickness. Interestingly, while the insertion efficiency for the largest QDs (11 nm) was 7.1% in the native membrane lacking PC24, it increased to 76.5% when inserted into the ‘thick’ membrane containing [PC24] = 25 mol%. For other NPs, we saw an increase from 46% to 70% for the 5 nm QDs, 40% to 48% for the 7 nm QDs, and 40% to 50% for the 5x13 nm NRs.

We further evaluated the degree of insertion for the 11 nm QDs by localizing the position of QDs relative to the SUVs’ membranes using a home-written software, as described in the SI. Figure 3D shows a histogram of these distances: positive distances represent locations in the outer SUV leaflet and outwards from center of the SUV, and negative distances represent locations in the inner SUV leaflet and inwards toward the center of the SUV. The histogram shows that these large NPs are most likely to be located near the center of the membrane. Figures 3(A-C) show three examples of analyzed FOVs, representing QDs that are located inward the SUV, the bilayer midline, and outward the SUV, respectively.

**Figure 3.**
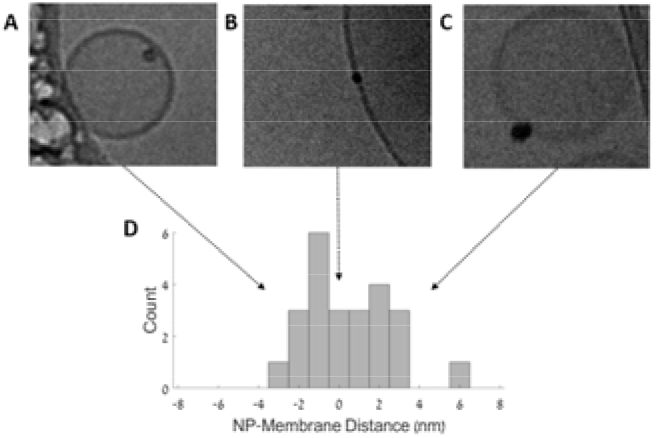
(a)-(c) Cryo-EM images showing 11 nm QDs inserted into thick SUVs. (d) Position distribution of QDs interacting with thick membrane SUVs. Only QDs that were in the vicinity of a SUV, but not in the vicinity of other QDs or more than one SUV, were counted (24 QDs in total).

We next performed molecular dynamics simulations to simulate the spherical QDs in their membrane-inserted states and characterize the role of PC24 in modulating lipid organization and structural properties to promote insertion efficiency. Due to the large size of these systems, we performed coarse-grained molecular dynamics simulations using the MARTINI^41^ force field, version 2.3P^42,43^, with the refined polarizable water model^44^, see Methods section for molecular dynamics simulations. Details on the CG models, simulation parameters, and simulation convergence are included in the SI (Tables S3-S5 and Figures S4-S6). Since the same trends were observed across the three QDs, we only show data for the intermediate 7 nm QD here; data for the 5 nm and 11 nm QDs are included in the SI (Figures S7-S13).

Figure 4 shows snapshots of final configurations of the 7 nm QDs in the native and thick membranes. The accompanying histograms show the volumetric density of headgroups (lipids/nm3) for each lipid species normalized by the volumetric density of the same headgroups in a bilayer without the inserted QD. These histograms provide information on the enrichment of each species compared to the bulk bilayer. Several observations are apparent from these data. First, comparing the POPC histograms at radial distances far from the QD reveals no significant increase in the average bilayer thickness in the thick membrane compared to the native membrane, likely because the fraction of PC24 (25 mol%) is small compared to POPC (45 mol%). Second, lipids extract from the membrane and onto the surface of the QD to shield the hydrophobic QD surface and ligands from water. At equilibrium, the inserted QDs are fully enveloped by lipids, in agreement with similar reports in the literature^45,46^. This behavior leads to substantial deformation of the extracted lipids due to the high curvature of the NP surface, which is expected to be unfavorable. Third, the simulation snapshots demonstrate the formation of bundles of hydrophobic ligands with voids between bundles^47^; these voids are filled with lipid tails (not visible in Figure 4 but shown in SI Figures S14-S16).

**Figure 4.** Simulation snapshots and 2D histograms of lipid headgroups for the 7 nm QD. Snapshots show the side view of the inserted QD with all water molecules, counterions, and half of the lipids in the *y*-axis with respect to the center of mass of the QD removed for visual purposes. The native membrane is shown at top and thick membrane at bottom. For the 2D histograms, *Z* is the distance along the *z*-axis and *D_r_* is the radial distance from the center of the QD in the *xy*-plane. The lipid headgroups selected for analysis were the phosphate group for POPC and PC24, choline group for DOTAP, and hydroxyl group for CHOL; these were binned radially in 0.375 nm wide by 0.375 nm tall bins and normalized by the volume of each bin and the number of configurations sampled. The color bar represents the fractional enrichment defined as the volumetric density of the lipid headgroups (lipids/nm^3^) normalized by the volumetric density of the lipid headgroups in the corresponding pure bilayer system.

A key distinction between the native and thick membranes is that PC24 is enriched in the region near the QD in the thick membranes (observed from the dark color for PC24 in Figure 4). This preferential enrichment results in an interfacial region around the QD that has a different lipid composition than the bulk bilayer. We further quantified enrichment by computing the area density of lipids (lipids/nm2) at fixed radial distances from the center of the QD. Figure 5A and Figure 5B reveal peaks in these area densities that corroborate lipid enrichment and depletion at defined radial distances. Specifically, in both the native and thick membranes the POPC and DOTAP densities are maximized at a radial distance 5-7 nm from the center of the QD, which we define as the QD-membrane interfacial region. We also observed that in thick membranes the area density for POPC and DOTAP decreases in this interfacial region while the area density of PC24 increases. Both observations are in agreement with their corresponding 2D histograms in Figure 4 and indicate that PC24 preferentially accumulates in the QD-membrane interfacial region, primarily displacing POPC.

**Figure 5.**
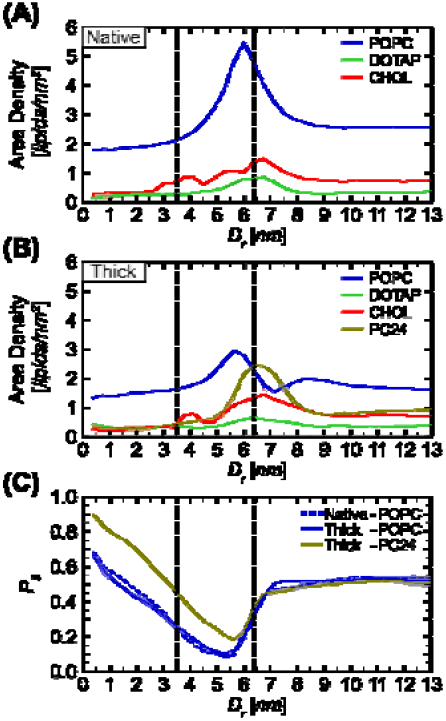
Lipid properties computed for the 7 nm QD. *D_r_* is the radial distance from the center of the QD in the *xy*-plane and parameters are calculated in radial bins with a width of 0.375 nm. The lipid headgroups selected for analysis were the phosphate group for POPC and PC24, choline group for DOTAP, and hydroxyl group for CHOL Lipid area density (lipids/nm^2^) for each type of lipid in the native **(A)** and the thick **(B)** membranes. **(C)** Lipid tail order parameter, computed for POPC and PC24 in both native and thick membranes. The left vertical dashed line corresponds to the QD radius, and the right vertical dashed line is the radial distance corresponding to the maximum area density of PC24. The error was calculated as the standard deviation between two replicas.

We hypothesized that the preferential enrichment of PC24 lipids in the QD-membrane interfacial region could be stabilizing QD insertion by reducing the extent of bilayer disruption. To test this hypothesis, we used MDTraj^48^ (v1.9.7) to compute the *P*_2_ lipid tail order parameter of the lipid acyl chain tails^49^, which is defined as:

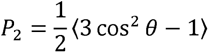

where *θ* denotes the angle between the vector normal to the bilayer and the vector from the second hydrophobic bead to the first hydrophobic bead bonded to the glycerol bead of the lipid tail. This parameter measures the structural orientation between consecutive tail atoms with respect to the bilayer normal; aligned lipid tails increase tail order. Figure 5C compares values of *P*_2_ for lipids near the QD to values of *P*_2_ to lipids far from the QD in both the native and thick membranes to quantify QD-driven membrane disruption (indicated by lower values of *P*_2_). Figure 5C shows that compared to its bulk value in the native membrane, *P*_2_ substantially decreases for the POPC lipids in the QD-membrane interfacial region where the area density of POPC reaches a maximum (Figure 5A). This observation is consistent with the energetically unfavorable disruption of lipid order in this interfacial region due to the presence of the QD. *P*_2_ increases for lipids extracted onto the QD, although this largely reflects the geometry of the curved QD surface. Similar behavior is observed for POPC in the thick membrane. Conversely, P_2_ decreases to a lesser degree for the long-chain PC24 lipids in the QD-membrane interfacial region, indicating less disruption of these lipids compared to the shorter-chain POPC lipids. Together, the data in Figure 5 indicate that in the thick membrane PC24 lipids preferentially accumulate in the QD-membrane interfacial region to reduce overall membrane disruption compared to the native membrane.

A second distinction between the native and thick membranes relates to the voids between ligand bundles observable in Figure 4; these voids are filled with lipid tails to minimize contact between hydrophobic ligand backbones and water. In the thick membrane, PC24 lipids preferentially partition into these void regions and exclude POPC lipids (SI Figures S14-S16), similar to the exclusion of POPC lipids from the QD-membrane interfacial region. These results suggest that the PC24 tails can more favorably partition into the void spaces given their longer length than the POPC tails, providing another mechanism to minimize overall bilayer disruption in the thick membranes.

## Discussion

Previous works have shown that only very small QDs (< 4 nm)^27–29^ can be incorporated into SUV membranes during their formation using electro-swelling, sonication and other detergent removal protocols^37^. Here, we used a different approach that relied on a previously published protocol for membrane incorporation of small hydrophobic drug molecules^33^. In this approach, a dialysis-based detergent removal step was performed during the preparation of the vesicles. This improved protocol was sufficient for the insertion of larger (> 5 nm) particles into SUVs with membranes composed of POPC and CHOL, although at low efficiency. The addition of 25 mol% of a long-chain phospholipid, PC24, significantly improved the insertion of large particles into SUV membranes, and facilitated the insertion of large QDs (∼11 nm) at high efficiency. Combining both approaches, NPs as large as 11 nm QDs and 5x13 nm NRs were successfully inserted into SUVs.

Coarse-grained molecular simulations suggest two key factors for why long-chain lipids facilitate the insertion of large CdSe/ZnS QDs coated with octadecylamine ligands, as schematically illustrated in Figure 6. First, long-chain lipids preferentially accumulate in the interfacial region at the QD-membrane interface, displacing POPC. The thicker interfacial region results in less disruption of the bilayer compared to the native membrane, which is energetically favorable and thermodynamically stabilizes QD insertion. PC24 thus acts similarly to a linactant^50^, a molecule that decreases the line tension between lipid domains. In this case, the line tension at the interface between lipids on the QD and lipids on the bulk bilayer is lowered, potentially increasing insertion efficiency for large NPs. Second, PC24 preferentially fills voids in the ligand monolayer that arise due to the formation of ligand bundles, again displacing POPC. The longer PC24 tails can more effectively pack against the long octadecylamine ligand backbones to again minimize bilayer disruption. These factors point to PC24-induced changes to lipid structural organization as facilitating QD insertion, even without substantial changes in overall membrane thickness.

**Figure 6.**
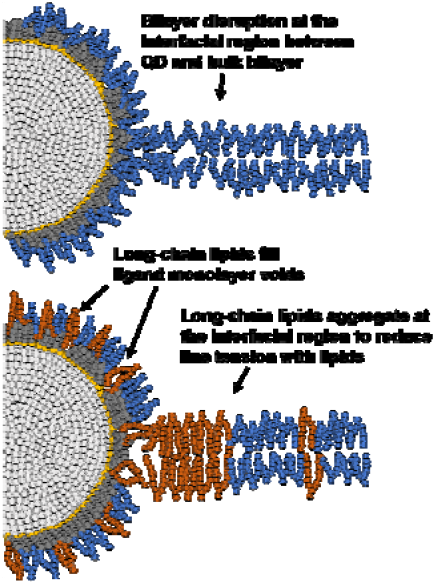
Schematic illustrating the long-chain lipid aggregation and increased lipid tail order in the interfacial region between the QD and the bulk bilayer (shaded region). Long-chain lipids (PC24) are colored orange. Medium-chain lipids (POPC and DOTAP) are colored blue. The QD shell is colored light gray, the amine group of the ligands are colored yellow, and the ligand hydrophobic groups are colored gray.

## Summary

In summary, we developed a lipid composition and SUV preparation protocol that promotes the integration of large inorganic hydrophobic particles into membranes. It is well-accepted that membrane thickness varies with acyl chain length and the presence of cholesterol. We found that adding long-chain phospholipids (C24:1) to the vesicle formulation is crucial for the incorporation of NPs with diameters much larger than those reported for typical lipid compositions. Molecular dynamics simulations indicate that the long-chain lipids act like linactants that stabilize membrane-inserted QDs. Although the main obstacle for NPs incorporation into SUVs is their size, the effect of the hydrophobic capping ligand layer and the particle geometry on insertion into SUVs is a matter of future study. These SUVs could be made fusogenic to allow the delivery of NPs into cellular membranes. The technology developed here could be applied to diagnostic technologies, therapeutic applications, and as a research tool for studying membrane properties. The computational approach in this work can also be potentially applied to other classes of nanomaterials and more biologically relevant membrane models to analyze insertion efficiency. By enabling these predictions, we can explore a wider range of materials as biomedical agents.

## Materials and Methods

### Chemicals

All chemicals of the highest purity available were purchased and were used without further purifications, buffer solutions, sodium cholate were purchased from Sigma-Aldrich. 1-palmitoyl-2-oleoyl-glycero-3-phosphocholine POPC, 1,2-dioleoyl-3-trimethylammonium-propane (chloride salt), DOTAP, Cholesterol and 1,2-dinervonoylsn-glycero-3-phosphocholine (PC24) were purchased from Avanti Polar Lipids. As-synthesized 5 nm QDs were purchased from Cytodiagnostics (Burlington, Canada), 7 and 11 nm QDs were purchased from ocean nanotech(San Diego, USA).

### Type II NRs synthesis CdSe/CdS

ZnSe/CdS core−shell NRs were synthesized following a previously published procedure with some modifications^20^. Then NRs were precipitated three times with 600 ul of toluene:methanol (1:1; v:v) solution. The NRs solution was then dissolved in CHCl3in order to remove excess ligands.

### TEM imaging

Carbon type A grid (Ted Pella Inc., Redding, USA) was glow discharged with a EmiTech K100 machine (Emitech Group, France) then 3µl of NP was loaded on the grid. After 1 minute the sample was blotted and access material was removed then air dried. The sample was then inspected with a Tecnai G2 microscope (FEI, Teramo fisher, Massachusetts, USA) with an acceleration voltage of 120 kV. Images were taken using Digital Micrograph with a Mulitiscan Camera model 794 (Gatan, Inc., Pleasanton, United States) in different resolutions.

### Absorbance spectroscopy

The absorbance spectrums of NPs were obtained with Ocean Optics USB4000 spectrophotometer. The spectra of NPs were obtained at a scan rate of 1200 nm/min with 1 cm path length in a quartz cuvette.

### Preparation of NPs embedded SUVs

In a typical preparation of a SUV/NP constructs, 7 µmol of total lipids (Table 1) were mixed with NPs (10 µL of 5 µM stock solution in chloroform) and dissolved in 30 µl Methanol: Methylene chloride: Chloroform (1:1:1; v:v:v). Next, 13 mg of the detergent sodium cholate was added to the organic solution giving 50 mM final concentration. In the next step, the organic solutions were evaporated for 1 h at 45°C using a vacuum evaporator. Afterwards, the dry film was re-suspended with 0.5 ml of 20 mM phosphate buffer pH 7.2, giving final lipids concentration of 18 mM, and the solution was equilibrated at 45°C for 1 h.

Detergent was removed by controlled dialysis of the mixture against 3 of 20 mM phosphate buffer pH 7.2 (volume ratio = 1-1000) at 45 °C (using GeBAflex-tube 8KDa MWCO), and mixture was extruded through 100 nm pore membrane 11 times at 45 °C using the LiposoFast instrument (Avanti polar lipids, Inc.)

### Dynamic light scattering

Dynamic light scattering data were collected using the Zetasizer Nano ZS instrument (Malvern Instruments, Great Britain) operating at 25 °C with a laser wavelength λ of 620 nm and a scattering angel of θ 173°. Measurements were made using polystyrene low-volume microcuvettes (BrandTech Scientific, Inc. USA). Cuvettes were filled with 100 μL of Vesicles dispersion for DLS characterization. DLS data is shown in Figure S1.

### Cryo-EM imaging

For the cryo-EM measurement, 3µl of SUV/NP samples were loaded on a glow discharged (EmiTech K100 machine) Quantifoil grid or lacey grid Grids that were blotted and plunged into liquid ethane using a Gatan CP3 automated plunger and stored in liquid nitrogen until use. Frozen specimens (samples with vesicles embedded in vitreous ice) were transferred to Gatan 914 cryo-holder and maintained at temperatures below -176°C inside the microscope. Samples were inspected with a Tecnai G2 microscope with an acceleration voltage of 120 kV, which is equipped with a cryobox decontaminator. Images were taken using Digital Micrograph with a Mulitiscan Camera model 794 in different resolutions.

### Data Analysis for insertion efficiency

The efficiency of NP-membrane incorporation was evaluated by calculation of the fraction of NPs that inserted SUVs lipid membrane. Three different experiments were conducted for each condition for every NP. The insertion efficiency was calculated as # membrane inserted NPs / total # NPs in a frame and the complementary non-inserted NPs as # non-inserted NPs / total # NPs in a frame. Error bars were averaged from 3 experiments.

### Simulation methods

We used GROMACS 2021.5^51,52^ to perform all simulations using a leap-frog integrator with a timestep of 20 fs. The MARTINI^41^ force field, version 2.3P^42,43^, with the refined polarizable water model^44^, was used to model the interactions. Energy minimization, equilibration, and production simulation parameters were the same for all systems. Energy minimization was performed for 5,000 steps or until the maximum force was below 1,000 kJ/mol·nm. System equilibration and production runs were performed in the *NPT* ensemble. The temperature was controlled at 300 K using a velocity-rescale thermostat with a time constant of 1 ps. During equilibration runs, the pressure was controlled at 1 bar using the Berendsen barostat with semi-isotropic pressure coupling with a time constant of 5 ps and compressibility of 3×10^-4^ bar^-1^. A 5 ns equilibration was performed with a timestep of 10 fs, followed by a 15 ns equilibration with a timestep of 15 fs, and a final 25 ns equilibration with a timestep of 20 fs, for a total of 45 ns of equilibration. Production runs were performed using the same parameters as the equilibration runs with the exception of the barostat, which was switched to a Parrinello-Rahman barostat with a time constant of 12 ps. We used the Visual Molecular Dynamics (VMD)^53^ software, release 1.9.4a55, to generate all simulation snapshots.

### System setup for coarse-grained MD simulations

We modeled the 5, 7, and 11 nm spherical CdSe/ZnS QDs as hollow shells coated with hydrophobic octadecylamine ligands. We coated the QD surface at a grafting density of 3.9 ligands/nm^2^ based on literature of similar CdSe/ZnS QDs^54,55,56^. Details on the coarse-grained models are included in the SI (Figure S4 and Table S3).

Lipids were modeled according to their standard topologies in MARTINI with PC24 modeled as DNPC. The MARTINI topology for DOTAP was added to this script. All bilayer lipid mixtures were generated using the *insane.py* script^57^. Pure (*i.e.*, without inserted QDs) native and thick bilayers were simulated in 15 nm × 15 nm × 10 nm simulation boxes for 100 ns to quantify their bulk properties; the last 50 ns were used for analyses (Figure S6). For the QD/bilayer systems, the QD was placed in the middle of the bilayer and solvated with polarizable water. We neutralized the system by adding a number of Cl^-^ ions equal to the number of DOTAP lipids using the *gmx genion* tool. The box size was 36 nm × 36 nm × 16 nm for the 5 nm QD, 38 nm × 38 nm ×16 nm for the 7 nm QD, and 42 nm × 42 nm × 20 nm for the 11 nm QD. We specified an initial area per lipid of 0.7 nm^2^/lipid to ensure system stability. The total number of molecules for all systems are listed in Table S4 and Table S5.

We performed production simulations of 200 ns for the systems with the 5 nm and 7 nm QDs and 400 ns for the system with the 11 nm QD, sampling every 0.2 ns. System convergence was assessed by plotting the area density (lipids/nm^2^) of all lipids throughout the production run (Figure S5). As the QD size increases, longer equilibration times were required. The last 100 ns of simulation time for all QD/bilayer systems were deemed as converged and used for subsequent analyses.

## Acknowledgements

We thank Prof. Dan Oron and Dr. Gaoling Yang for providing the NR sample. This work has received funding from the European Research Council (ERC) under the European Union’s Horizon 2020 research and innovation program under grant agreement No. 669941 and ERC-POC grant agreement No. 779896, by the BER program of the Department of Energy Office of Science grant DE-SC0020338, by the STROBE National Science Foundation Science & Technology Center, Grant No. DMR-1548924, by the Israel Science Foundation grant # 813/19, and by the Bar-Ilan Research & Development Co, the Israel Innovation Authority, Grant No. 63392. C.A.H.Z and R.C.V. acknowledge the support provided by the Graduate Engineering Research Scholars – Advanced Opportunity Fellowship from the University of Wisconsin – Madison, the National Institute of General Medical Sciences’ Chemistry-Biology Interface Training Program (5T32GM008505-29) from the University of Wisconsin – Madison, and from the National Science Foundation CAREER Award No. DMR-2044997. This work used the Advanced Cyberinfrastructure Coordination Ecosystem: Services & Support (ACCESS), which is supported by the National Science Foundation under Grant No. 2138307.

## Supporting Information Available

DLS data, NPs characteristics, SUV/NP preparations components, parameters of the coarse-grained model, number of molecules for each QD/bilayer system and simulation snapshots.

## Notes

### Competing Interest Statement

The authors have declared no competing interest.

### Summary of Updates

The manuscript has undergone major revisions and MD modeling was added to this work.

