## Supplemental Information for "Long-chain lipids facilitate insertion of large nanoparticles into membranes of small unilamellar vesicles"

**Coarse-grained model for CdSe/ZnS quantum dots**

**Table S1:** Characteristics of NPs used for the SUV/NP samples in this study.

| NP | Composition | Ligands | Diameter (nm) | Emission wavelength (nm) | Extinction coefficient (M^-1^CM^-1^) |
| --- | --- | --- | --- | --- | --- |
| Trilite525 | CdSeS/ZnS | Hydrophobic | 4.42± 1.07 | 525 | 570,000 |
| QSP620 | CdSe/ZnS | ODA | 6.91± 0.91 | 620 | 640,000 |
| QSP600 | CdSe/ZnS | ODA | 11.06± 1.99 | 600 | 900,000 |
| NRs | CdSe/ZnS | TOPO | 5x13± 2.19 | 600 | 1,630,000 |

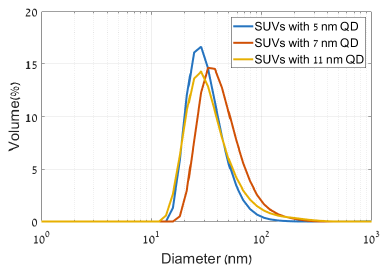

**Figure S1:** DLS size distribution by volume. For all the three SUVs one single peak were observed by intensity. Mean size by intensity is 60.09 ± 32.82 nm for the 5nm, 87.49 ± 47.28 nm for the 7nm and 102.5 ± 68.12nm for the 11nm.

**Table S2:** Components of native and thick membranes for SUV/NP preparations.

| Membrane Composition | | Native Membrane (%) | Thick Membrane (%) |
| --- | --- | --- | --- |
| POPC | 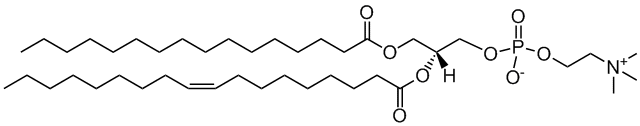 | 70 | 45 |
| CHOL | 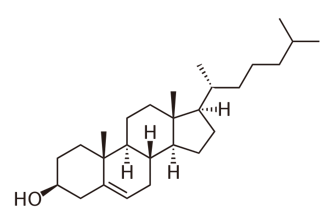 | 20 | 20 |
| DOTAP | 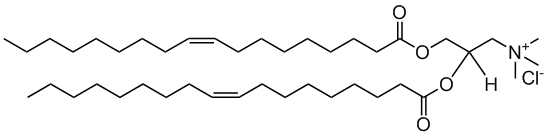 | 10 | 10 |
| PC24 | 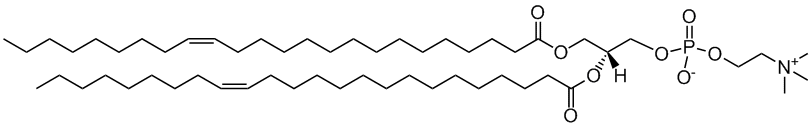 | 0 | 25 |

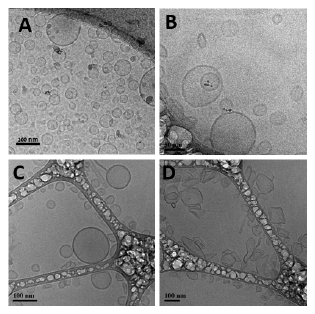

**Figure S2:** representative cryo-EM images showing examples of SUVs with increasing PC24% in the lipid composition. (a) 0% PC24, (b) 25% PC24, (c) 75% PC24, (d) 100% PC24.

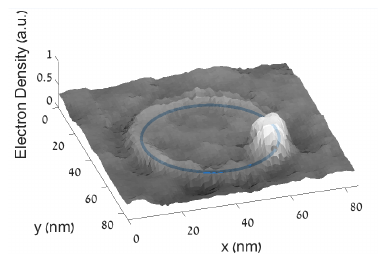

**Figure S3:** Representation of the center of mass position of QDs(marked with a blue dot) calculated by averaging the weighted position of each pixel in the cryo-EM image, with the weight being the image gray values, which corresponds to the local electron density. By fitting the membrane curve (marked with a blue circle) in the vicinity of the particle to a circle, the distance between the QD’s position and membrane was defined as the minimal distance between the membrane curve and the QD’s center of mass.

We performed coarse-grained molecular dynamics simulations using version 2.3P^1,2^ of the MARTINI^3^ force field with the polarizable water model^4^ to model all system components. CdSe/ZnS quantum dots (QDs) are mainly synthesized in organic environments with the aid of hydrophobic ligands^5,6^ and prior studies have found that ZnS surfaces are mildly hydrophobic.^7^ Therefore, we modeled the QD core as a spherical hollow shell using moderately hydrophobic C3 beads. The number of shell beads for each QD was calculated by dividing the surface area of the QD by the area occupied by each shell bead. The surface area of the shell was calculated as $A=4\pi r^{2}$ using the corresponding radius of each QD plus 0.24 nm to account for the radius of the shell beads. The area of a single shell bead was calculated as $A=\pi r^{2}$ using an interatomic diameter of 0.47 nm scaled by 0.9. Scaling down the area of each shell bead was required to minimize voids between them and was crucial to prevent ligands from unphysically partitioning towards the inside of the shell. The shell beads were bonded to their nearest neighbors within 0.59 nm for the 5 and 7 nm QDs and within 0.94 nm for the 11 nm QD. These cutoffs were optimized by scaling the interatomic diameter by a factor that yielded a stable and rigid shell. Shell beads were bonded together at an equilibrium length corresponding to the nearest distances with a harmonic potential of 5,000 kJ/mol·nm^2^ for rigidity. The structure and topology files for the coarse-grained QDs were generated using in-house Python scripts.

Each octadecylamine ligand was modeled using hydrophobic apolar beads as shown in Figure S4. The N(CH_2_)_3_ headgroup was modeled as a C5 bead, the (CH_2_)_4_ groups were modeled as C1 beads, the remaining (CH_2_)_3_ was modeled as a SC1 bead. Alkanes are commonly modeled using apolar beads using the MARTINI force field. We modeled the propylamine as a less hydrophobic bead to retain the hydrophobic nature of the QDs. Every C5 bead was bonded to its closest C3 bead with a harmonic potential of 5,000 kJ/mol·nm^2^. We obtained the bonded parameters for the ligand beads through a bottom-up parameterization by reproducing atomistic data and analyzing the bond length and bond angle distributions. The details of this bottom-up parameterization will be included in a forthcoming manuscript. The bonded parameters are listed in Table S3.

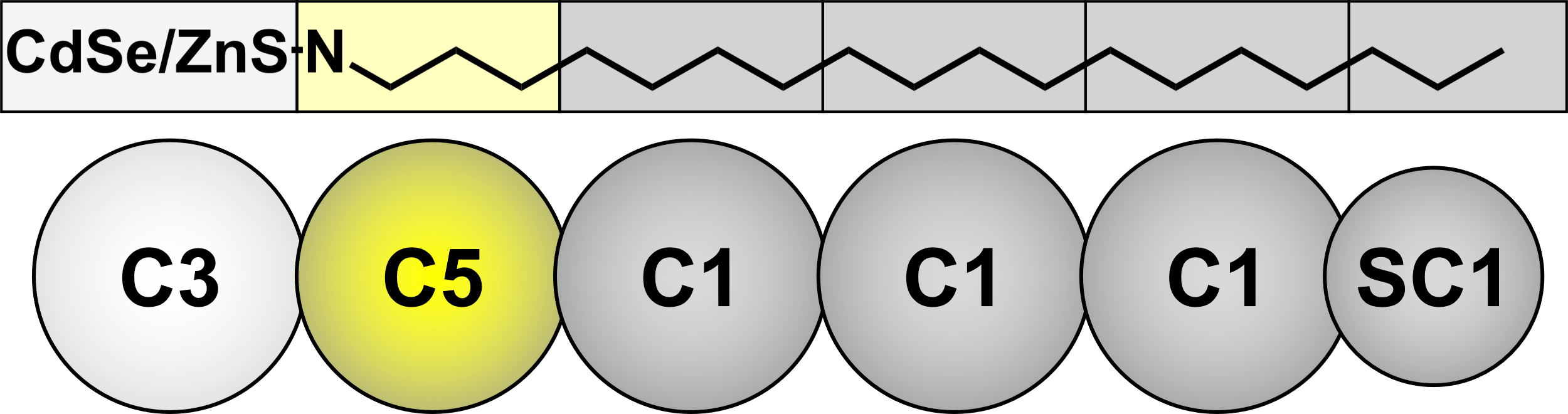

**Figure S4:** Chemical structure of the octadecylamine ligand compared to its coarse-grained model labeled according to the chosen MARTINI topologies.

**Table S3:** Bonded parameters for the coarse-grained model of the octadecylamine ligand.

| Beads | Length | Force Constant | Beads | Angle | Force Constant |
| --- | --- | --- | --- | --- | --- |
|  | [nm] | [kJ/mol·nm^2^] |  | [degrees] | [kJ/mol] |
| C5-C1 | 0.52 | 12,200 | C5-C1-C1 | 165 | 538 |
| C1-C1 | 0.51 | Constraint | C1-C1-C1 | 175 | 1,149 |
| C1-C1 | 0.51 | Constraint | C1-C1-SC1 | 172 | 264 |
| C1-SC1 | 0.44 | 9,300 |  |  |  |

**Table S4:** Total number of molecules for the pure bilayers.

| Replica #1 | Native | Thick |
| --- | --- | --- |
| POPC | 452 | 290 |
| DOTAP | 64 | 64 |
| CHOL | 128 | 128 |
| PC24 | 0 | 162 |
| Polarizable Water | 10,380 | 9,579 |
| Cl^-^ | 64 | 64 |

**Table S5:** Total number of molecules for each QD/bilayer system.

| Replica #1 | 5 nm | | 7 nm | | 11 nm | |
| --- | --- | --- | --- | --- | --- | --- |
|  | Native | Thick | Native | Thick | Native | Thick |
| CdSe/ZnS | 558 | | 1,095 | | 2,704 | |
| Octadecylamine | 367 | | 685 | | 1,614 | |
| POPC | 2,450 | 1,575 | 2,629 | 1,690 | 3,109 | 1,998 |
| DOTAP | 349 | 349 | 374 | 374 | 443 | 443 |
| CHOL | 699 | 699 | 750 | 750 | 887 | 887 |
| PC24 | 0 | 875 | 0 | 938 | 0 | 1109 |
| Polarizable Water | 120,519 | 116,163 | 132,243 | 127,489 | 210,654 | 205,364 |
| Cl^-^ | 349 | 349 | 374 | 374 | 443 | 443 |
| Replica #2 | 5 nm | | 7 nm | | 11 nm | |
|  | Native | Thick | Native | Thick | Native | Thick |
| CdSe/ZnS | 558 | | 1,095 | | 2,704 | |
| Octadecylamine | 367 | | 685 | | 1,614 | |
| POPC | 2,450 | 1,575 | 2,629 | 1,690 | 3,109 | 1,998 |
| DOTAP | 349 | 349 | 374 | 374 | 443 | 443 |
| CHOL | 699 | 699 | 750 | 750 | 887 | 887 |
| PC24 | 0 | 875 | 0 | 938 | 0 | 1,109 |
| Polarizable Water | 120,542 | 116,198 | 132,301 | 127,465 | 210,687 | 205,335 |
| Cl^-^ | 349 | 349 | 374 | 374 | 443 | 443 |

**Convergence of molecular dynamics simulations**

To assess the convergence of each system, we plotted the area density (lipids/nm^2^) of all lipids throughout the production run as shown in Figure S5. As the QD size increases, longer equilibration times were required. The last 100 ns of simulation time for all QD/bilayer systems were deemed as converged and used for subsequent analyses.

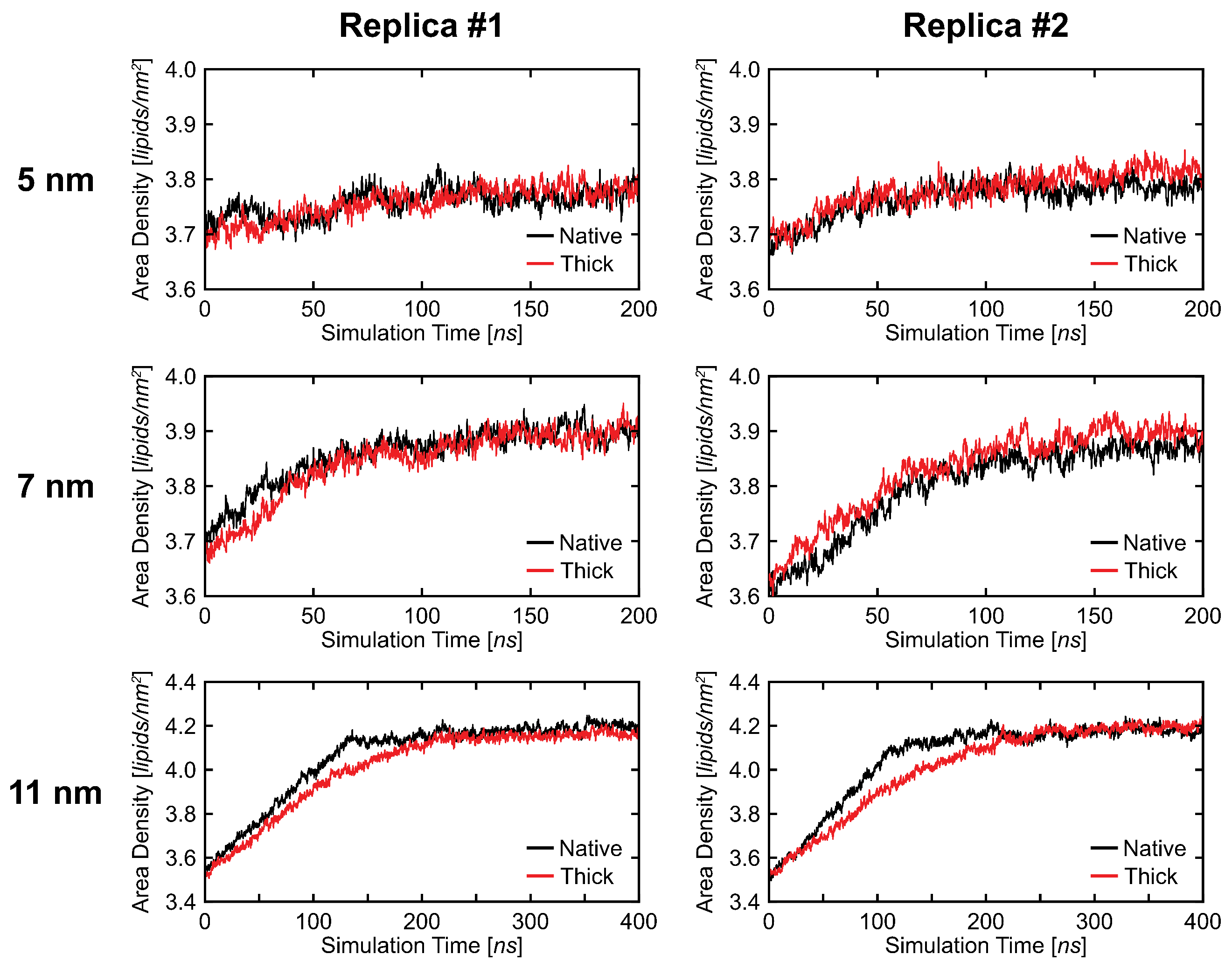

**Figure S5:** Area density plots as convergence metric. The area density was calculated for every frame sampled using the headgroup of each lipid as reference (phosphate group for POPC and PC24, choline group for DOTAP, hydroxyl group for CHOL). Lipids from both leaflets were included in the calculation to maintain consistency with the analyses performed.

**Supporting data**

The time-averaged thickness of the pure bilayers was calculated using the *z*-coordinate of the phosphate group for POPC and PC24, and the choline group for DOTAP. Sampling was performed every 1 ns of the last 50 ns of the production run. The thickness was then computed as the difference between the average of *z­*-coordinates of upper leaflet lipids and the average of ­*z*-coordinates of lower leaflet lipids divided by the 50 ns sampled. The time-averaged thicknesses were 4.09 nm and 4.50 nm for the pure native and thick bilayers, respectively, indicating an ~10% increase in thickness upon incorporation of PC24. Simulation snapshots are shown in Figure S6.

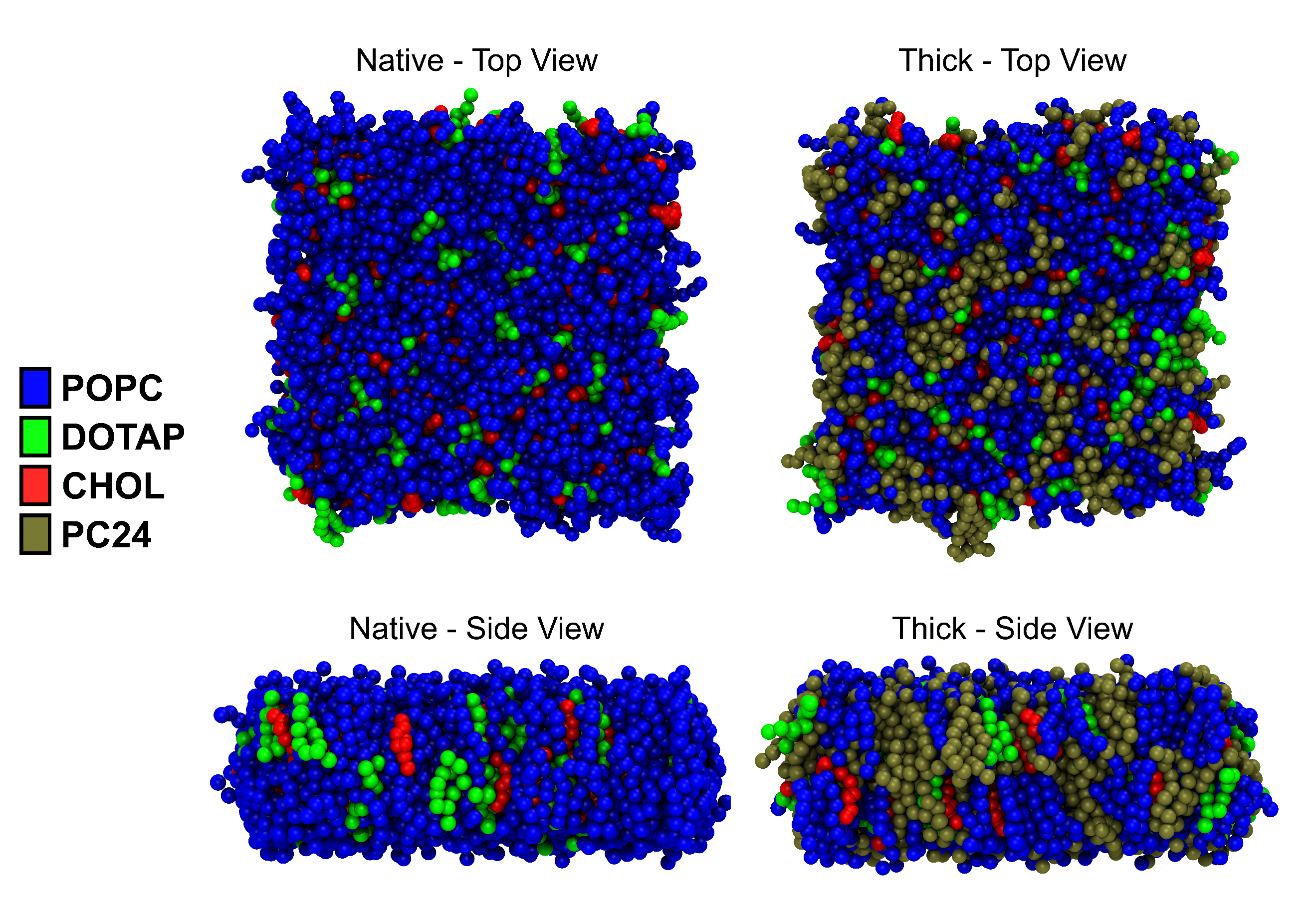

**Figure S6:** Simulation snapshots of the final configuration from a 100 ns production run. Top and side views for both native and thick membranes are shown.

The 2D histogram data for the 5 and 11 nm QDs (two replicas), and for the second replica of the 7 nm QD are included below. Figures from the main text are not included.

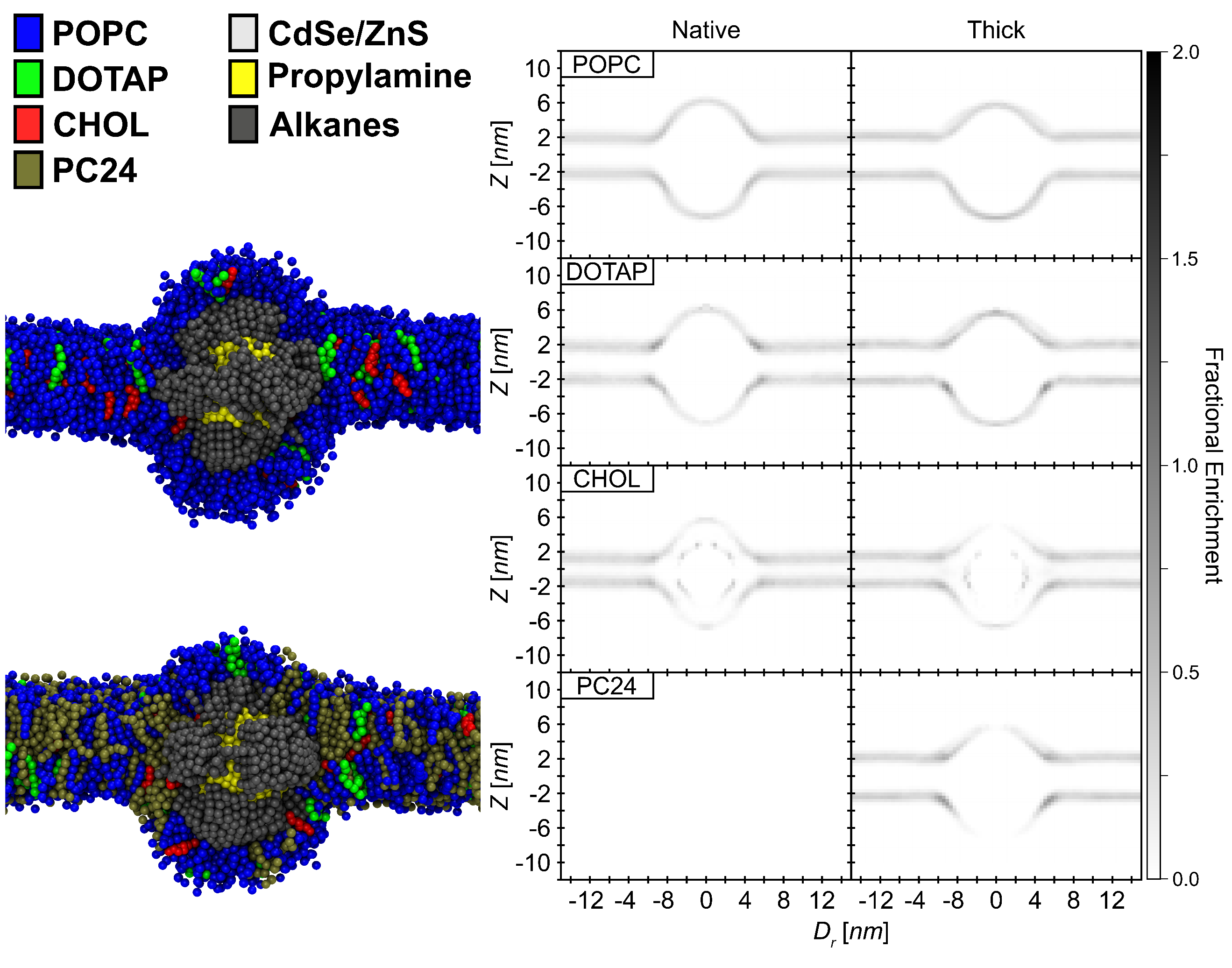

**Figure S7:** Simulation snapshots and 2D histograms of lipid headgroups for the first replica of the 5 nm QD showing the side view of the inserted QD. For the simulation snapshots, all water molecules, counterions, and half of the lipids in the *y*-axis with respect to the center of mass of the QD were removed for visual purposes. For the 2D histograms, *Z* is the *z*-axis and *D_r_* is the radial distance from the center of the QD; values were adjusted to the center of the QD, and negative *D*_r_ values were plotted for visual purposes. The lipid headgroups were binned radially in 0.375 nm wide by 0.375 nm tall bins and normalized by the volume of each bin and the number of configurations sampled. The lipid headgroups selected for analysis were the phosphate group for POPC and PC24, choline group for DOTAP, and hydroxyl group for CHOL. The color bar represents the fractional enrichment defined as the volumetric density of the lipid headgroups (lipids/nm^3^) normalized by the volumetric density of the lipid headgroups in the corresponding pure bilayer system.

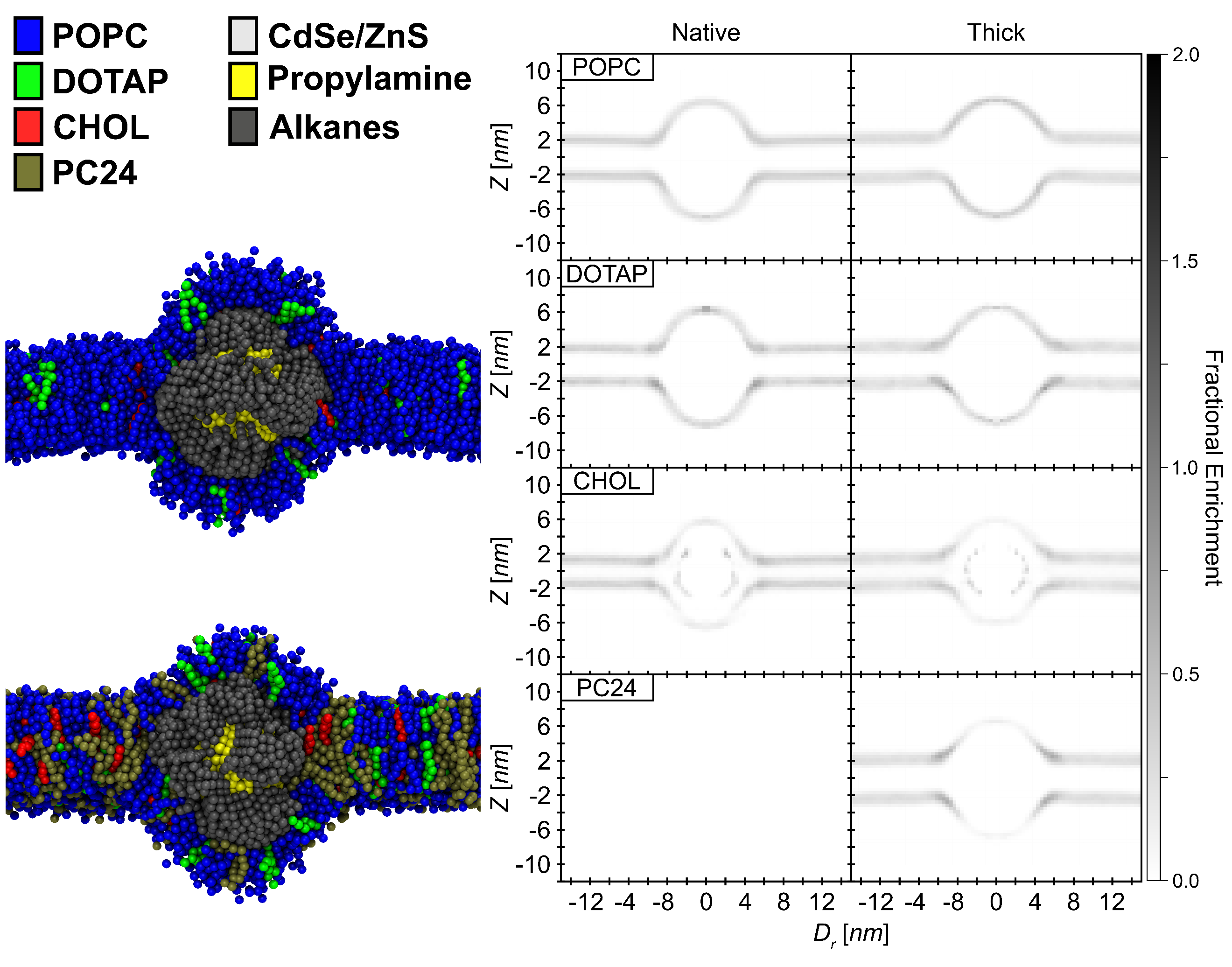

**Figure S8:** Simulation snapshots and 2D histograms of lipid headgroups for the second replica of the 5 nm QD showing the side view of the inserted QD. For the simulation snapshots, all water molecules, counterions, and half of the lipids in the *y*-axis with respect to the center of mass of the QD were removed for visual purposes For the 2D histograms, *Z* is the *z*-axis and *D_r_* is the radial distance from the center of the QD; values were adjusted to the center of the QD, and negative *D*_r_ values were plotted for visual purposes. The lipid headgroups were binned radially in 0.375 nm wide by 0.375 nm tall bins and normalized by the volume of each bin and the number of configurations sampled. The lipid headgroups selected for analysis were the phosphate group for POPC and PC24, choline group for DOTAP, and hydroxyl group for CHOL. The color bar represents the fractional enrichment defined as the volumetric density of the lipid headgroups (lipids/nm^3^) normalized by the volumetric density of the lipid headgroups in the corresponding pure bilayer system.

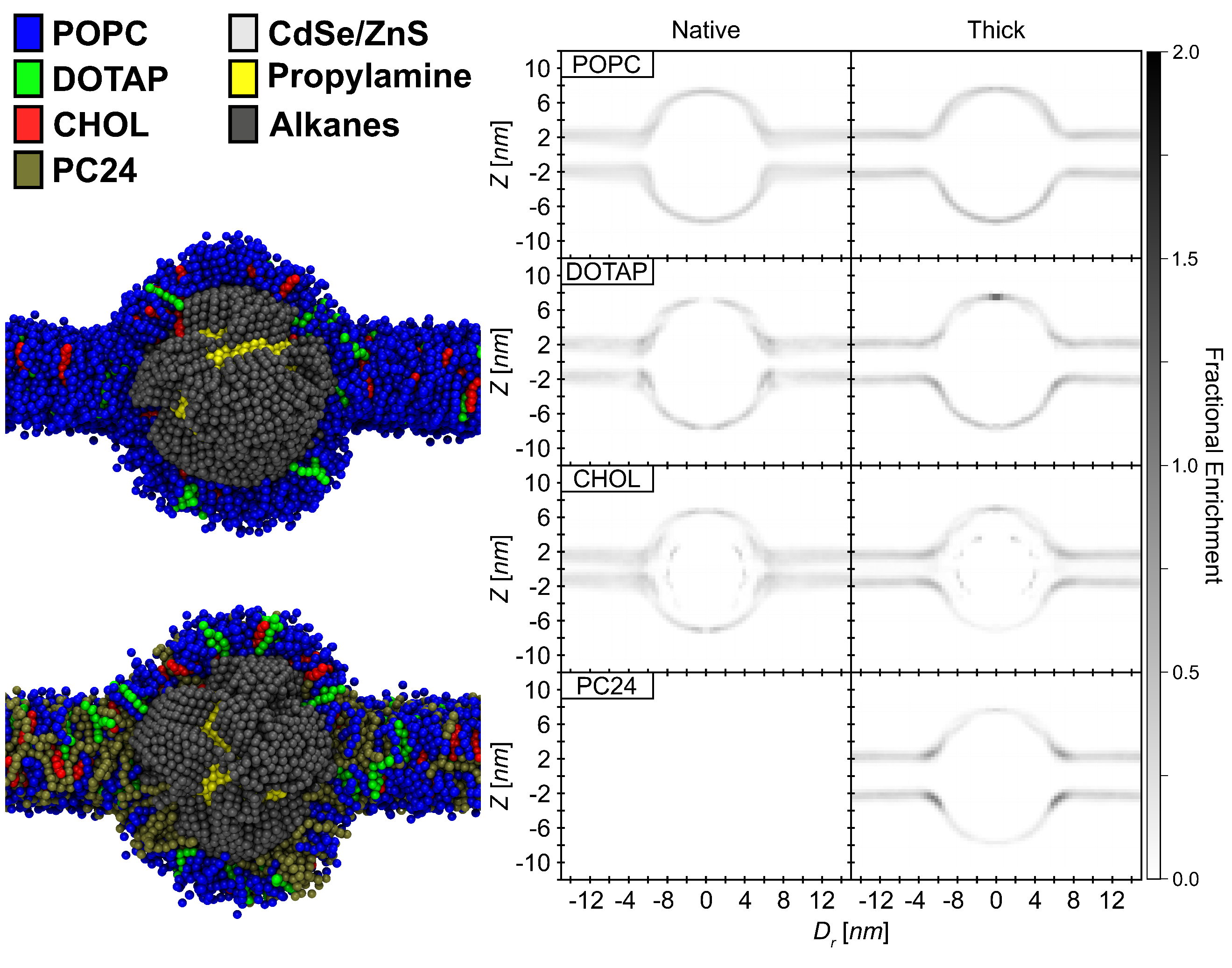

**Figure S9:** Simulation snapshots and 2D histograms of lipid headgroups for the second replica of the 7 nm QD showing the side view of the inserted QD. For the simulation snapshots, all water molecules, counterions, and half of the lipids in the *y*-axis with respect to the center of mass of the QD were removed for visual purposes. For the 2D histograms, *Z* is the *z*-axis and *D_r_* is the radial distance from the center of the QD; values were adjusted to the center of the QD, and negative *D*_r_ values were plotted for visual purposes. The lipid headgroups were binned radially in 0.375 nm wide by 0.375 nm tall bins and normalized by the volume of each bin and the number of configurations sampled. The lipid headgroups selected for analysis were the phosphate group for POPC and PC24, choline group for DOTAP, and hydroxyl group for CHOL. The color bar represents the fractional enrichment defined as the volumetric density of the lipid headgroups (lipids/nm^3^) normalized by the volumetric density of the lipid headgroups in the corresponding pure bilayer system.

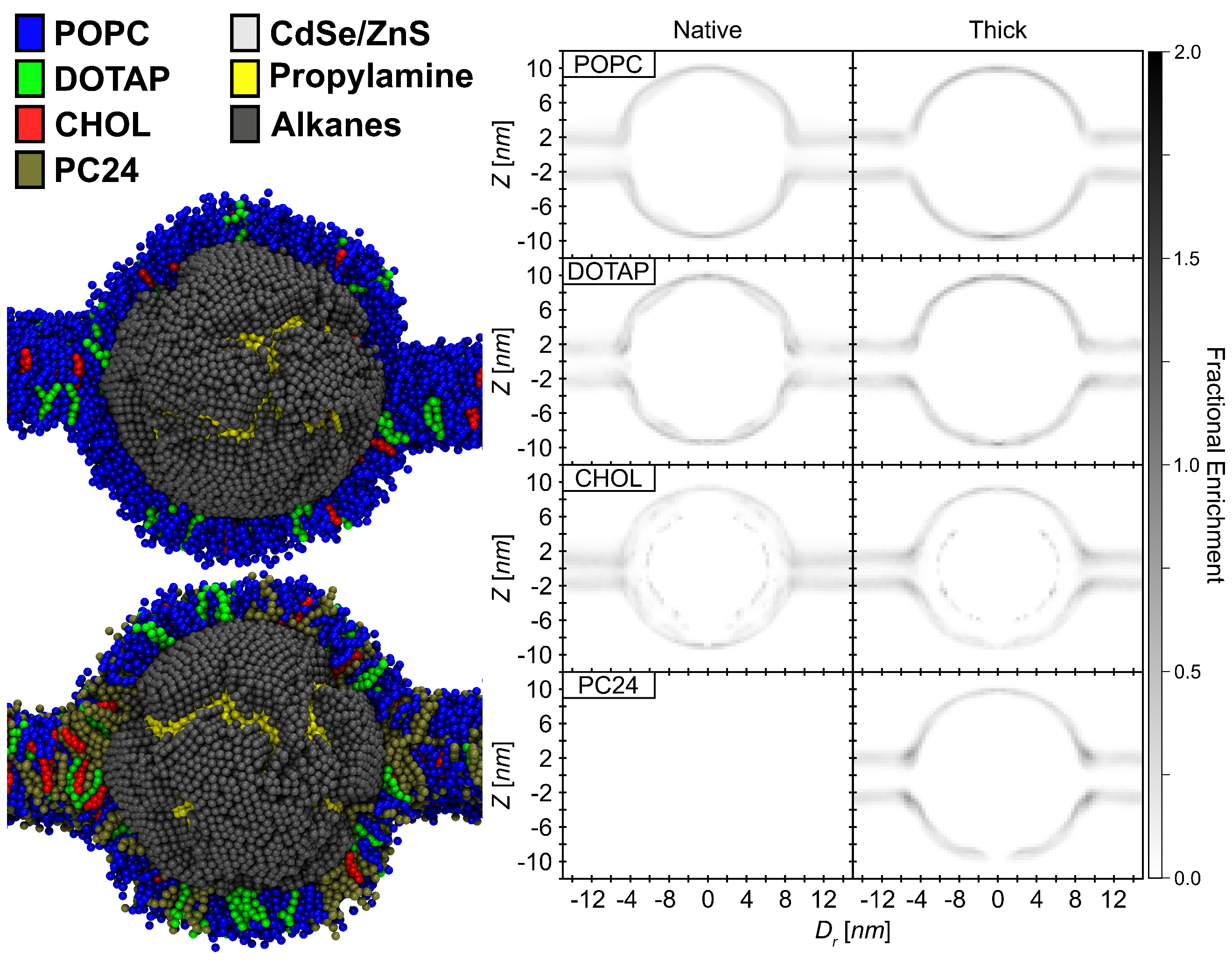

**Figure S10:** Simulation snapshots and 2D histograms of lipid headgroups for the first replica of the 11 nm QD showing the side view of the inserted QD. For the simulation snapshots, all water molecules, counterions, and half of the lipids in the *y*-axis with respect to the center of mass of the QD were removed for visual purposes. For the 2D histograms, *Z* is the *z*-axis and *D_r_* is the radial distance from the center of the QD; values were adjusted to the center of the QD, and negative *D*_r_ values were plotted for visual purposes. The lipid headgroups were binned radially in 0.375 nm wide by 0.375 nm tall bins and normalized by the volume of each bin and the number of configurations sampled. The lipid headgroups selected for analysis were the phosphate group for POPC and PC24, choline group for DOTAP, and hydroxyl group for CHOL. The color bar represents the fractional enrichment defined as the volumetric density of the lipid headgroups (lipids/nm^3^) normalized by the volumetric density of the lipid headgroups in the corresponding pure bilayer system.

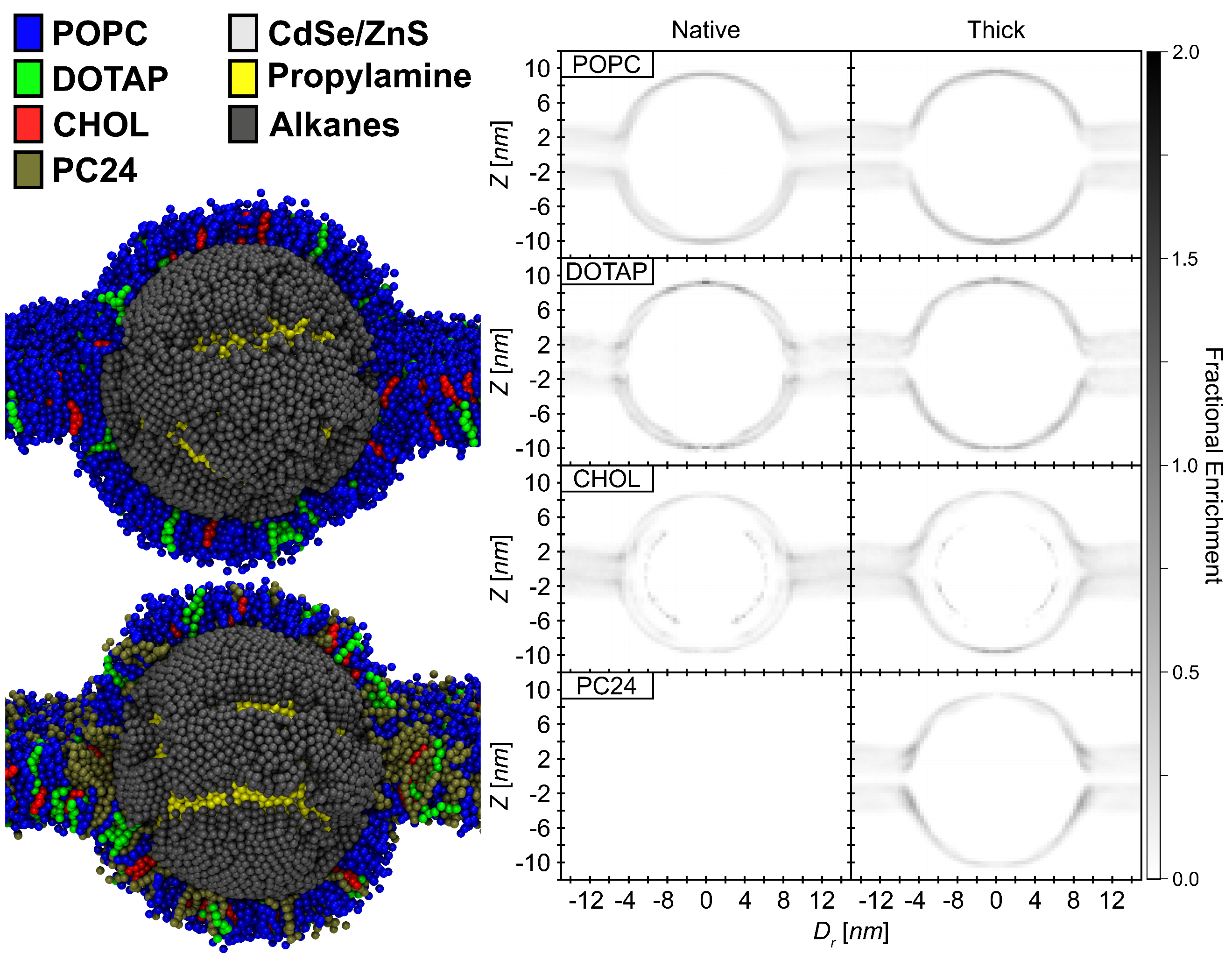

**Figure S11:** Simulation snapshots and 2D histograms of lipid headgroups for the second replica of the 11 nm QD showing the side view of the inserted QD. For the simulation snapshots, all water molecules, counterions, and half of the lipids in the *y*-axis with respect to the center of mass of the QD were removed for visual purposes For the 2D histograms, *Z* is the *z*-axis and *D_r_* is the radial distance from the center of the QD; values were adjusted to the center of the QD, and negative *D*_r_ values were plotted for visual purposes. The lipid headgroups were binned radially in 0.375 nm wide by 0.375 nm tall bins and normalized by the volume of each bin and the number of configurations sampled. The lipid headgroups selected for analysis were the phosphate group for POPC and PC24, choline group for DOTAP, and hydroxyl group for CHOL. The color bar represents the fractional enrichment defined as the volumetric density of the lipid headgroups (lipids/nm^3^) normalized by the volumetric density of the lipid headgroups in the corresponding pure bilayer system.

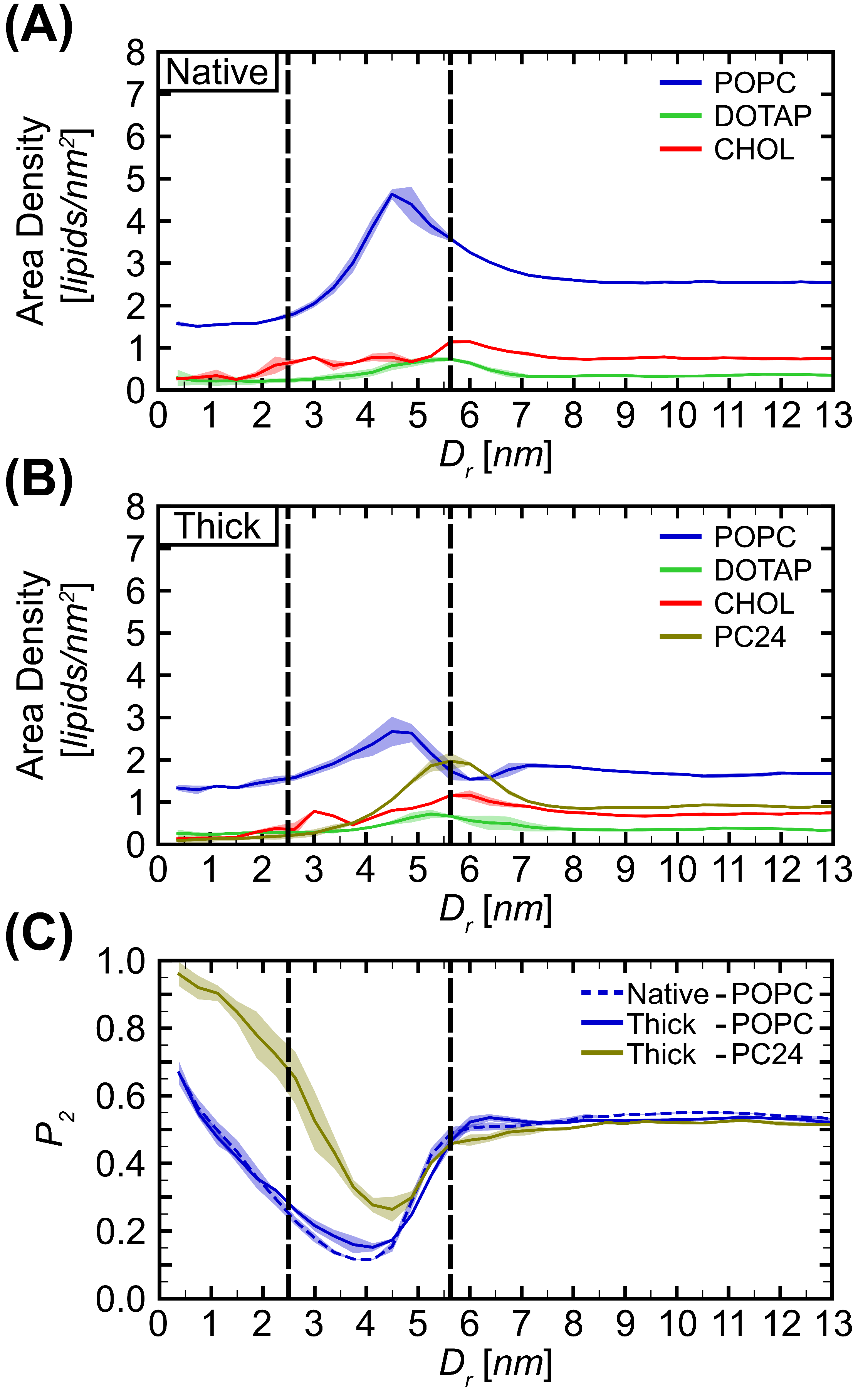

**Figure S12:** Lipid properties computed for the 5 nm QD. *D_r_* is the radial distance from the center of the QD in the *xy*-plane and parameters are calculated in radial bins with a width of 0.375 nm. The lipid headgroups selected for analysis were the phosphate group for POPC and PC24, choline group for DOTAP, and hydroxyl group for CHOL Lipid area density (lipids/nm^2^) for each type of lipid in the native **(A)** and the thick **(B)** membranes. **(C)** Lipid tail order parameter, $P_{2}$, computed for POPC and PC24 in both native and thick membranes. The left vertical dashed line corresponds to the QD radius, and the right vertical dashed line is the radial distance corresponding to the maximum area density of PC24. The error was calculated as the standard deviation between two replicas.

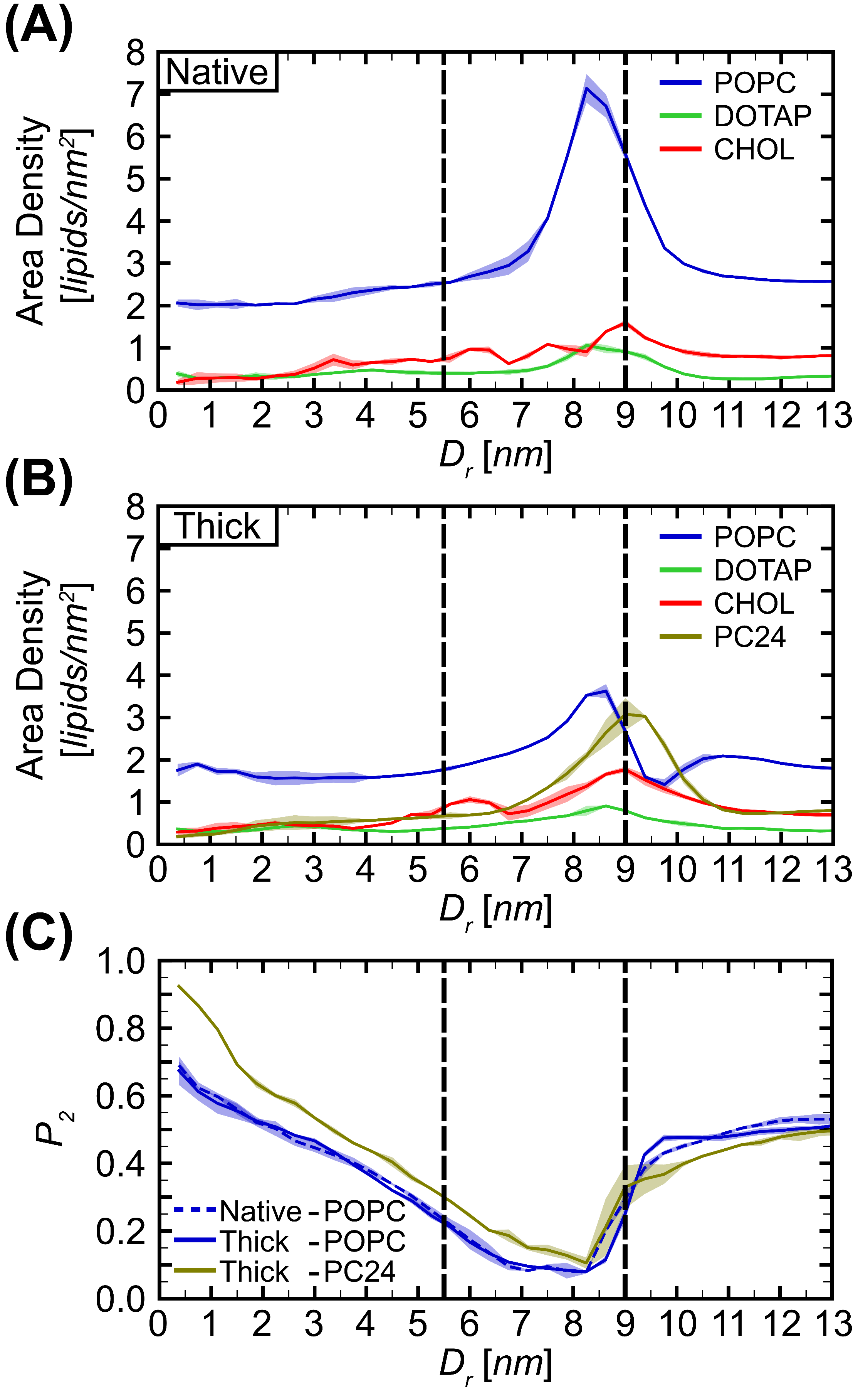

**Figure S13:** Lipid properties computed for the 11 nm QD. *D_r_* is the radial distance from the center of the QD in the *xy*-plane and parameters are calculated in radial bins with a width of 0.375 nm. The lipid headgroups selected for analysis were the phosphate group for POPC and PC24, choline group for DOTAP, and hydroxyl group for CHOL Lipid area density (lipids/nm^2^) for each type of lipid in the native **(A)** and the thick **(B)** membranes. **(C)** Lipid tail order parameter, $P_{2}$, computed for POPC and PC24 in both native and thick membranes. The left vertical dashed line corresponds to the QD radius, and the right vertical dashed line is the radial distance corresponding to the maximum area density of PC24. The error was calculated as the standard deviation between two replicas.

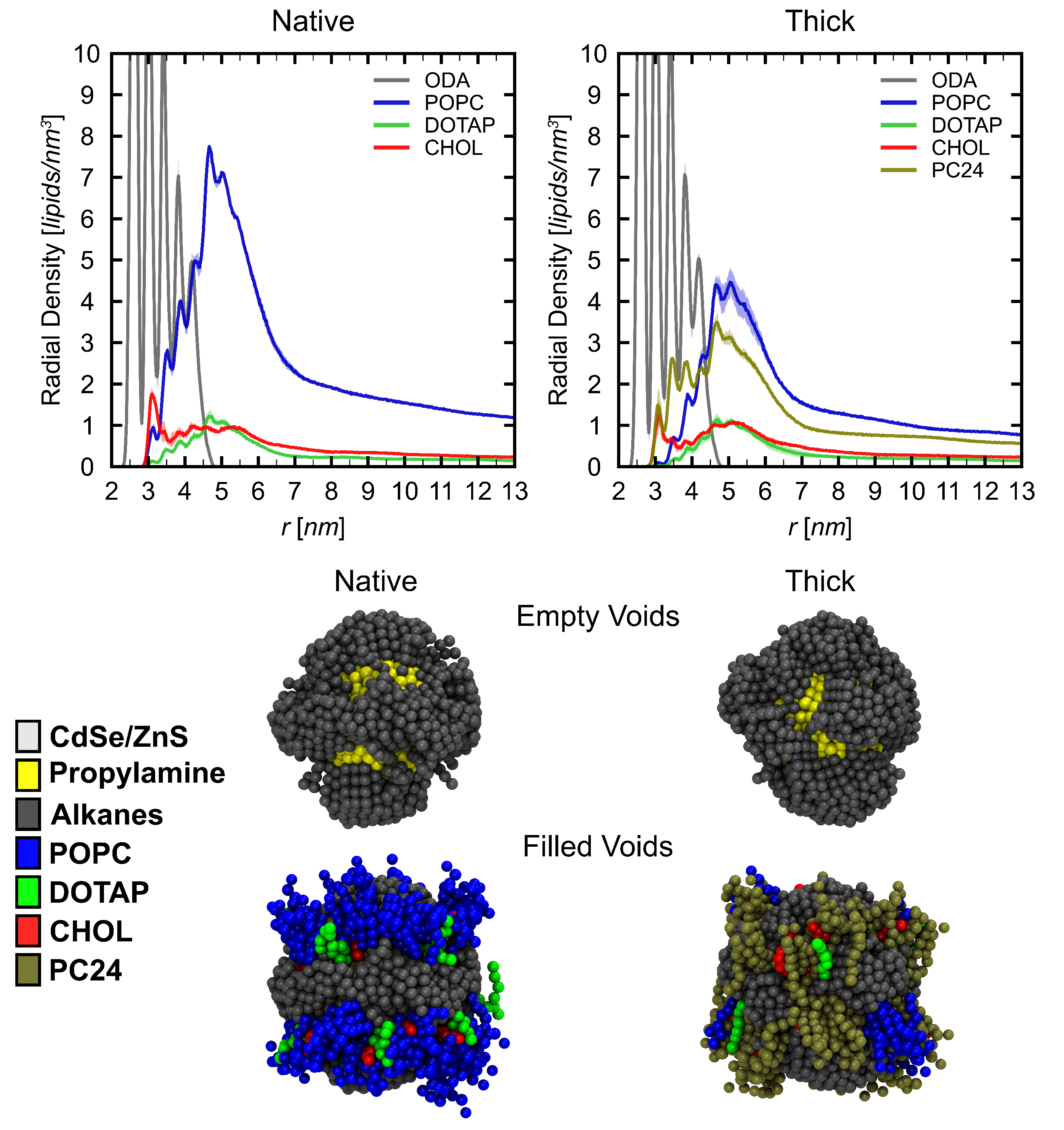

**Figure S14:** Lipid distribution around the 5 nm QD. Radial distribution functions were computed as a function of the distance from the center of the QD (in 3D, as opposed to in the *xy*-plane) and normalized to yield number densities using the *gmx rdf* tool with a bin size 0.02 nm. The error was calculated as the standard deviation between two replicas. Only lipids with at least one bead within 1 nm of the S1 beads are shown on the simulation snapshots.

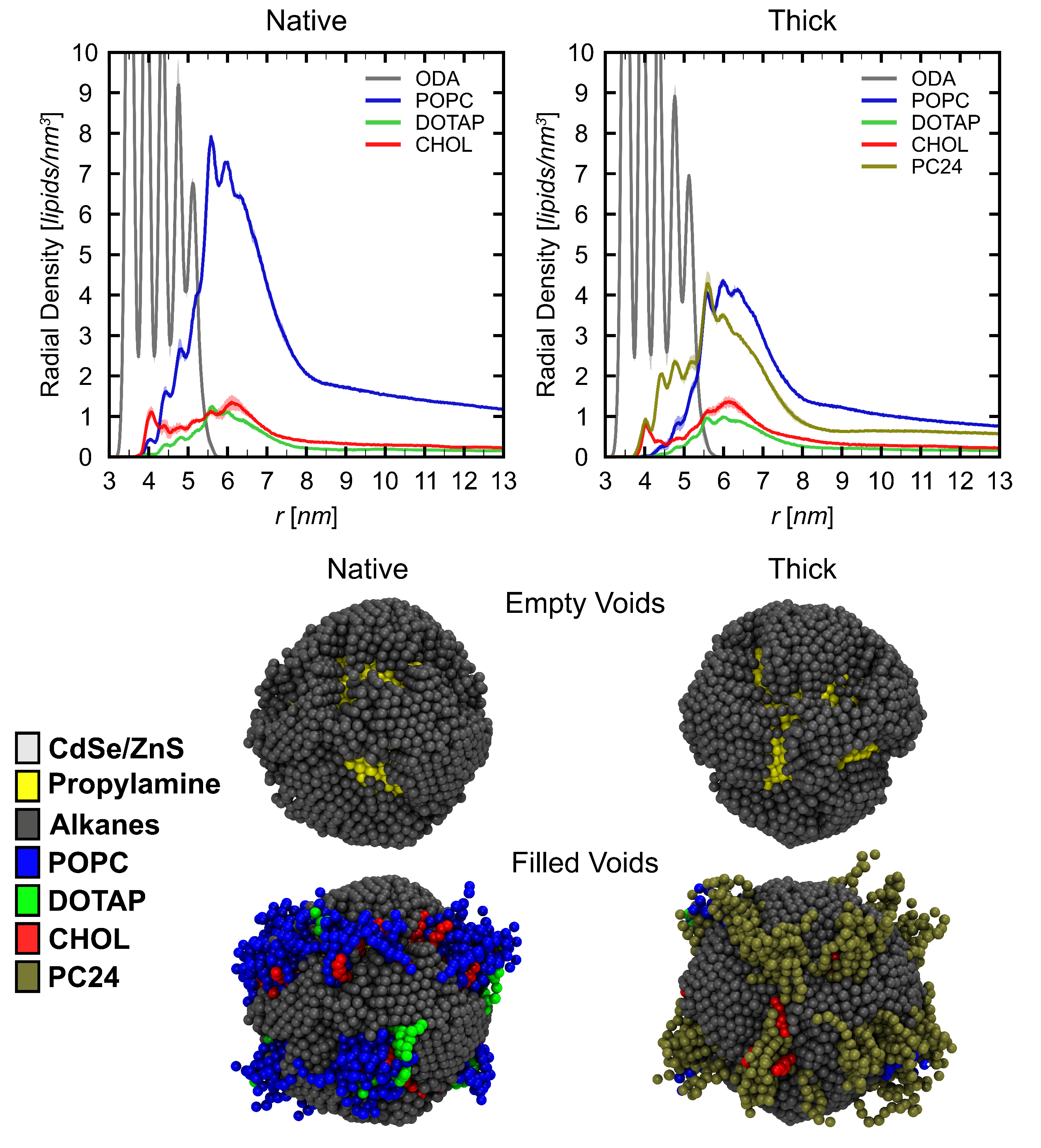

**Figure S15:** Lipid distribution around the 7 nm QD. Radial distribution functions were computed as a function of the distance from the center of the QD (in 3D, as opposed to in the *xy*-plane) and normalized to yield number densities using the *gmx rdf* tool with a bin size 0.02 nm. The error was calculated as the standard deviation between two replicas. Only lipids with at least one bead within 1 nm of the S1 beads are shown on the simulation snapshots.

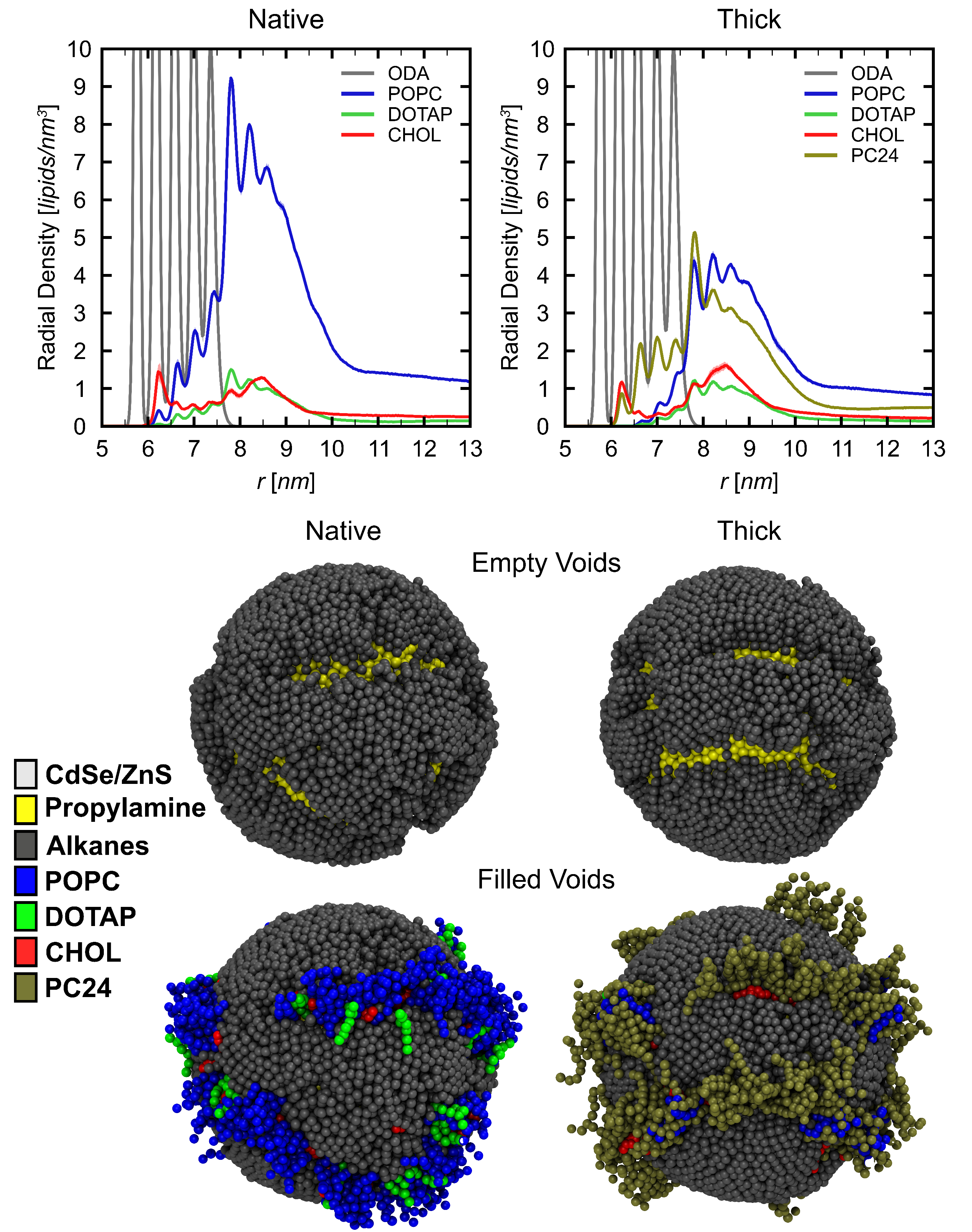

**Figure S16:** Lipid distribution around the 11 nm QD. Radial distribution functions were computed as a function of the distance from the center of the QD (in 3D, as opposed to in the *xy*-plane) and normalized to yield number densities using the *gmx rdf* tool with a bin size 0.02 nm. The error was calculated as the standard deviation between two replicas. Only lipids with at least one bead within 1 nm of the S1 beads are shown on the simulation snapshots.
